# Modelling mental disorders in zebrafish. Neurexins severely modulate anxiety, social behaviours and aggression

**DOI:** 10.1101/2025.11.24.690344

**Authors:** Q Nguyen, F. Guo, B. Mowry, J. Das, J. Giacomotto

**Affiliations:** Institute for Biomedicine and Glycomics, Griffith University, Brisbane, QLD 4111, Australia; School of Environment and Science Griffith University, Brisbane, QLD 4111, Australia; Queensland Centre for Mental Health Research, The Park, Centre for Mental Health, Wacol, QLD, 4076, Australia; Queensland Brain Institute, The University of Queensland, Brisbane, Queensland, 4072, Australia; Thomson Institute, National PTSD Research Centre, University of the Sunshine Coast, Birtinya, QLD, Australia

## Abstract

Can we reproduce mental disorders in zebrafish? To explore this question, we manipulated the neurexin gene family. Neurexin (*nrxn*) genes encode synaptic cell-adhesion molecules that have been repetitively associated with neurodevelopmental and mental disorders. While the zebrafish animal model offers tremendous advantages for dissecting neural development/function, no robust *nrxn* zebrafish models are currently available.

In this study, we generated the first collection of zebrafish knockout lines for each *nrxn* gene, with mutations ranging from transmembrane domain- to full genomic locus-deletions. Surprisingly, all homozygous lines developed normally, presenting no gross neurodevelopmental or obvious early behavioural abnormalities. However, this absence of early phenotypes translated into profound, paralog-specific behavioural alterations emerging during juvenile-to-adult stages.

All neurexin knockouts affected mating behaviour, complicating the generation and maintenance of homozygous lines. Except for this shared behavioural alteration, *nrxn1*, but not *nrxn2* or *nrxn3*, led to marked changes in social behaviour and aggression. In contrast, *nrxn2* mutants exhibited severe anxiety-like behaviours, including bottom-dwelling and repetitive freezing/seizure events. Strikingly, *nrxn1* full-locus deletion mutants showed opposing behaviour, spending most of their time near the surface. The two also displayed opposite responses to open/closed field transitions; confinement alleviated *nrxn2* anxiety but enhanced *nrxn1* surface-dwelling. Meanwhile, *nrxn3* mutants behaved normally in all our initial tests.

In summary, our study introduces a complete set of zebrafish mutants covering the whole *nrxn* gene family, presenting striking juvenile/adult behavioural alterations despite the absence of noticeable early defects; echoing the delayed onset of human psychiatric disorders such as schizophrenia. This work demonstrates the value of zebrafish to study mental disorders and unlock a novel platform to unravel the pathogenic contribution of neurexin and associated subtle neurodevelopmental changes/timing that drive the emergence of mental illnesses.

## INTRODUCTION

Neurexins, *NRXN1*, *NRXN2* and *NRXN3* in humans, code for transmembrane cell-adhesion molecules essential for synaptic formation, maintenance and signalling^1,2^. These genes have repeatedly been associated with diverse mental and neurodevelopmental disorders such as Schizophrenia (SCZ), Autism (ASD), bipolar disorder (BD) and even recently with Parkinson (PD)^1–6^. A vast number of rodent models have been developed over the last two decades, which have significantly contributed to gaining tremendous insights into the biology of this complex gene family^2,4^. Despite this tremendous effort, their pathogenic role and association with the aforementioned diseases remain unclear. Expanding the tools available to broaden our ability to interrogate these genes is still essential. Surprisingly, while the zebrafish seems to be a model of choice to tackle the function of these genes, especially in investigating their role in the developing brain, there is currently a lack of available robust models for research.

Indeed, complementing their rodent counterparts, the zebrafish offers a unique set of tools for studying neurodevelopment and greater versatility for drug discovery^7^. They also exhibit extra-utero development and a very convenient optical transparency, enabling direct visualisation and manipulation of their brain even during the very early developing stages. In addition, the emergence of optogenetics and calcium imaging techniques with this organism offers unique ways to strengthen the ongoing effort aiming at understanding the normal and pathogenic role of this gene family in the developing and mature brain^8,9^. Despite these tremendous advantages, no robust zebrafish neurexin models are currently available for research. This small animal presents highly conserved neurexin orthologs, which have been ancestrally duplicated, neurexin 1 (*nrxn1a* and *nrxn1b*), neurexin 2 (*nrxn2a* and *nrxn2b*) and neurexin 3 (*nrxn3a* and *nrxn3b*)^10^.

These genes are transcribed into hundreds of isoforms due to multiple promoters and complex, still poorly understood, alternative splicing^11^. This complexity makes the generation of robust loss-of-function (LOF) mutations challenging. Traditional small indels - generating premature stop-codons and truncated proteins- are at high risk of being genetically compensated by *de-novo* and/or cryptic splicing. Let alone genetic compensation, it is not straightforward to target all isoforms at once due to the multiple promoters leading to a variety of α-long and β-short isoforms. Although aberrant splicing and compensatory mechanisms may contribute to disease pathogenesis and warrant closer investigation^12^, we aimed here at first generating a set of robust zebrafish knockout lines properly missing the associated gene/protein function, and to subsequently assess their effect on zebrafish development and behaviour, as well as the utility of this model organism for studying mental disorders.

Here, we are using an innovative CRISPR/Cas9 strategy to generate a complete set of zebrafish mutants covering the whole neurexin gene family and conduct a first systematic phenotyping comparison of each homozygous mutant line from embryonic to adult stage. We show that loss of these genes does not markedly affect early development or basic motor function, but results in striking and paralog-specific behavioural alterations in adulthood. Given that most mental health disorders (such as schizophrenia) emerge without detectable developmental or anatomical/structural abnormalities, while being accepted to be neurodevelopmental in origin, the presented data suggest that the zebrafish could be a valuable model to uncover pathogenic mechanisms in play prior to the symptoms. Our study also validates the zebrafish as a powerful tool to uncover the complex role of neurexins in brain development and function.

## MATERIAL AND METHODS

### Zebrafish maintenance

Adult zebrafish and embryos were maintained by standard protocols approved by the University of Queensland and Griffith University Animal Ethics Committee. Ethics approval AE213_18/AE213_18 and GRIDD/11/22/AEC. Wildtype and all lines used in these studies are in the AB background.

### *Neurexin* mutants’ generation and maintenance

All mutants presented in this study were generated following the protocol described in Tromp *et al.* 2021^13^. CRISPR guide RNAs and targeting sequences are listed in **Supplemental Table S1**. Genotyping of the heterozygotes and homozygotes was performed using the primer sets presented in **Fig. 1** and **Supplemental Table S2**. PCRs were carried out on genomic DNA extracted from dechorionated embryos or fin clips using the REDExtract-N-Amp Tissue PCR Kit (Sigma-Aldrich, Cat. XNAT-10RXN), according to the manufacturer’s instructions. Annotated sequences including sgRNA target sites, primer binding sites, and generated deletions are provided in **Supplemental File S1**.

**Figure 1.**
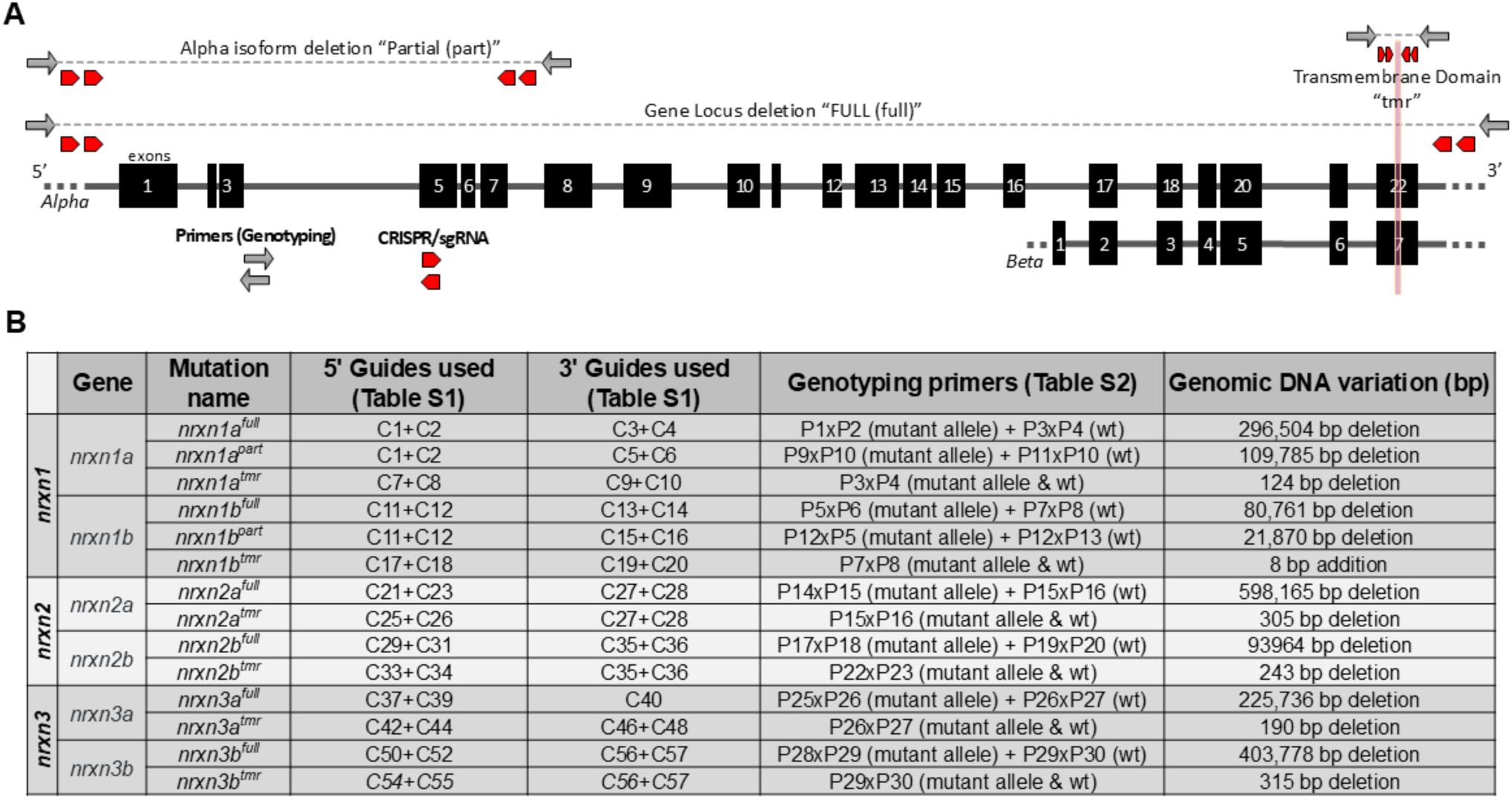
Schematic representation and details of the mutations generated across the six zebrafish neurexin genes. A, Schematic genomic organisation of the zebrafish neurexin loci showing the relative positions of the CRISPR/Cas9 single-guide RNAs (sgRNAs; arrowheads) used to generate the mutations/deletions described in this study. mutations/deletions presented in this study. Grey arrowheads indicate the approximate positions of the genotyping primers used for screening and maintenance of the different mutant lines. For the large deletions, the identification of the homozygous mutants required two sets of primers, with combination presented in the bottom panel. B, Summary of the different neurexin alleles/mutants generated, including corresponding i) sgRNA used (supplemental table S1), ii) genotyping primer pairs for identification and maintenance (supplemental table S2), and iii) genomic deletion sizes.

### Embryos and larvae measurements

Embryonic and larval development were analysed using a MVX10 Microscope (Olympus, 0.63x) equipped with a DP75 digital camera and controlled with CellSens image analysis software (Olympus). Body length was measured from calibrated images, defined as the distance from the anterior tip of the head to the posterior tip of the tail. Measurements were performed on 30 larvae per genotype at 3, 5, and 7 days post-fertilisation (dpf). Hatching dynamics were assessed across multiple developmental time points at 24, 32, 40, 48, 56, 64, 72, 80, 88, and 96 hours post-fertilisation (hpf). The number of hatched embryos was counted manually at each time point in multiple independent clutches for each mutant line and wild-type controls. These data were used to generate hatching curves and identify potential genotype-specific differences in developmental timing.

### Swimming-evoked response assays

Touch-evoked responses were assessed at 48 hpf in larvae from seven mutant lines and wild-type controls. Embryos were manually dechorionated at 24 hpf and maintained at 28.5 °C until testing. Prior to the assay, larvae were transferred to room temperature for one hour. Individual larvae (n = 30 per genotype) were placed into 90-mm Petri dishes containing E3 embryo medium. Mechanical stimuli were gently applied to the tail using a fine pestle tip. Each larvae received up to ten touches, with a successful escape response scored as ‘1’. Larval movements were recorded using the ZebraBox Revolution system (ViewPoint Life Sciences, France) until swimming ceased following stimulation. Swimming tracks were analysed in FIJI (ImageJ 1.8.0_322, 64-bit), and response rates were plotted as mean ± SEM using GraphPad Prism (version 9.0.0).

### Brain imaging and brain volume quantification

To assess brain morphology, we used an established transgenic line, *Tg(HuC:Kaede)*, expressing the fluorescent protein Kaede under the control of the HuC pan-neuronal promoter, generated through Tol2-mediated genomic integration^14^. Brain scans were acquired using an Olympus FV3000 laser scanning confocal microscope. Embryos were raised in E3 medium supplemented with 0.2mM 1-Phenyl-2-thiourea (PTU; final concentration) to block pigmentation. Larvae were scanned at 33, 57, 81, 105, 129, and 153 hpf at one-hour intervals using the 488nm excitation channel. Acquired z-stack images were processed and analysed using FIJI (ImageJ 1.8.0_322, 64-bit) and Imaris software (v. 10.2.0 Oxford Instruments, UK). Three-dimensional brain models were reconstructed using the surface-rendering function in Imaris, applying a uniform fluorescence-intensity threshold across all samples to standardise brain-volume quantification.

### Larvae swimming and behaviour analysis

Larval swimming behaviour and response to stress stimuli were assessed using the ZebraBox Revolution system (ViewPoint Life Sciences). Larvae were distributed in 24-well plates in triplicate, with each plate containing 12 control (AB background) and 12 mutant animals. The behavioural protocol consisted of a 32-minute recording, comprising four cycles of 4 minutes of light and 4 minutes of dark. During recording, repetitive 3-second light flashes (2400 lm) and 3-second acoustic vibration (250 Hz) were delivered every minute (independently offset by 30 seconds). Larvae were habituated in the dark for 5 minutes prior to recording. Data were analysed using GraphPad Prism (version 9.0.0).

### Adult behavioural studies

All adult behavioural experiments (described below) were performed using Zantiks LT automatic system (Zantiks Ltd., Cambridge, UK). All the tests were conducted between 11 am and 6 pm to minimise circadian disruption. The experimental room was maintained under constant temperature and illumination conditions. Mutant lines were tested in parallel with wild-type controls and statistical comparison was restricted to batches performed within the same timeframe.

### Novel tank diving test

The Novel Tank Diving test (also known as the Novel Tank Test, NTT) uses vertical distribution of zebrafish in a novel environment as a validated measure of ‘anxiety-like’ behaviours in adult zebrafish^15^, and was performed following established protocols^16^. Fish were acclimatised for at least 1 hour in the experimental room prior to behavioural testing. Each zebrafish was placed in a transparent tank of 125 x 155 (mm) dimensions (T-225 cell culture flask, supplied by Zantiks Ltd., Cambridge, UK). Fish behaviours were recorded for 15 min in lateral view and tracked via the Zantiks Software interface. For data analysis, the tank was virtually divided into top (T), middle (M), and bottom (B) zones. Total distance travelled, average time spent in each zone, and average entries into each zone of the tank were generated in real time using the Zantiks in-built software. Average velocity, latency to first entry into the top zone, number of freezing bouts and freezing percentage were subsequently quantified using the EthoVision XT software (Ver. 18.0.1803, Noldus, Netherlands).

### Mirror biting test

The mirror biting is a well-established fish paradigm^17,18^. The test was conducted in opaque tanks (20 × 14 × 14 cm) to minimise visual distractions from the external environment. A mirror insert was placed along one side of the tank, with two mirrors fitted at an angle to enhance the reflective surface^19^. The mirror zone was defined as the area within 3 cm of the mirror. Each fish was individually acclimated in the tank for 5 min in the dark. Following acclimation, illumination was restored, exposing the fish to the mirrors, and behaviour was recorded for 10 min. The total time spent within the mirror zone was calculated as the primary measure of mirror-directed behaviour.

### Social preference test

The social preference assay was designed to evaluate social interaction and preference in zebrafish^18,20^. An individual test fish was introduced into the central compartment of an opaque testing tank, with a group of same-strain conspecifics placed in an adjacent compartment at one end. The opposite end contained an empty compartment, serving as a neutral control. The central compartment was further virtually divided into 4 equal-sized subzones; the zone nearest to the friend fish was designated “social zone”, the 2 zones in the middle was designated “middle zone”, and the third zone was designated “empty zone”. Behavioural recordings started after an acclimation of 5 min. Videos were recorded from a top view of the tank for 10 min. The time spent in each zone was then calculated and social preference index was computed to quantify the degree of social interaction exhibited by the test fish.

### Closed Field Test

The principle of this assay is similar to the Novel Tank Test (NTT); however, it has been adapted in our lab to assess anxiety-like behaviour in a group of fish rather than in individuals. This assay is referred to as the Closed Field Test, as it was conducted within a confined environment inside the Zantiks LT system. This contrasts with the Open Field Test (described below), where the tank is placed in an open environment external to the Zantiks system, leaving the animal exposed to the surrounding space (illuminated and with a large field of view). The Closed Field Test was performed in a transparent rectangular tank (37 x 14 x 30, mm, length × width × height). A group of 10 fish was gently placed into the tank, and their behaviour was recorded for 15 min from lateral view. The experiment was repeated four times (n = 4 independent groups of 10 fish; 40 fish in total). The tank height was virtually divided into three horizontal zones: top (T), middle (M), and bottom (B). The number of fish in each zone was manually counted every 5 seconds using Adobe Premium Pro 2022 (Adobe Inc., California, USA). The shoal was recorded for 15 minutes, including a 5-minute acclimatisation and a 10-minute experimental recording used to manually plot zone distribution across time.

### Open Field Test

This assay used the same rectangular tank used in the Closed Field Test but installed in an open setting outside the Zantiks LT system, exposed to the surrounding space - free of an investigator and in dedicated illuminated room-. To ensure consistency, water depth was maintained at the same level as in the Closed Field Test. A light panel was positioned behind the tank to improve contrast and animal detection. Behavioural recordings were captured from the side using a Logitech Brio 4K Pro Webcam (Logitech International S.A., Lausanne, Switzerland) connected to the Zantiks Console control system. As in the Closed Field Test, each 10-fish group was gently netted into the tank, and their behaviour were recorded for 15 min (5 min acclimatisation), following the same analytical pipeline. Four groups (n = 4 independent groups of 10 fish; 40 fish in total) were tested per genotype.

### Swimming tunnel assay

This test was conducted using a swimming tunnel apparatus from Loligo® Systems (Madison, USA), with protocol modifications based on a previous study^21^. A group of 4 fish was placed into the glass chamber, and behaviour was recorded laterally using a “Logitech BRIO – Ultra HD Webcam”. Animals were first acclimated for 5 min in static water. Following acclimation, water velocity was increased stepwise by 10 cm/s every 5 min to induce flow-driven swimming behaviour. The time from the onset of flow to exhaustion was recorded, with exhaustion defined as the point at which the fish could no longer maintain position within the flow and were carried onto the collection mesh at the downstream end of the chamber. The moment a fish ceased active swimming was recorded as its time to exhaustion.

### Statistical analysis

Statistical analysis was performed using GraphPad Prism 9 software (GraphPad, La Jolla, CA, USA). Data are presented as mean ± SEM. Comparisons between groups were conducted using statistical tests indicated in the figure legends. In the figures, * indicates p < 0.05, ** indicates p < 0.01, *** indicates p < 0.001, and ns indicates no significance.

## RESULTS

### Zebrafish neurexin (*nrxn*) mutants’ generation

To generate LOF zebrafish *nrxn* mutants, we employed a CRISPR/Cas9 methodology previously established in our laboratory to facilitate the generation of targeted large deletions across the genome^13^. We applied this approach to generate mutants for all zebrafish *nrxn* genes, *e.g. nrxn1a*, *nrxn1b*, *nrxn2a*, *nrxn2b*, *nrxn3a*, and *nrxn3b*, and established double-mutant colonies for phenotypic investigations. We slightly modified the published technique/methodology to use only four guides (*e.g.* minus the described fifth guide targeting the gene tyrosinase as a selection marker); with two guides on each side of the deletion of interest as described in **Figure 1**. To avoid genetic compensation via alternative splicing as well as enable the study of non-coding regions across these genome loci, we aimed to remove i) the entire genomic regions, including all introns (deletions named “full” in this manuscript) or ii) each transmembrane region (located on the last exon of each gene; deletions named “tmr”). For *nrxn1*, we also generated an additional line with a deletion/mutation running from the α-isoform promoter to Exon 7 (named “part”), which does not overlap with the genomic region associated with the short *nrxn1*-isoforms, *nrxn1a-β* and *nrxn1b-β*. The details of these deletions are presented in **Figure 1**, with the corresponding annotated sequences provided in **Supplemental File S1**, and all guide RNAs and genotyping primers listed in **Supplemental Table S1** and **Supplemental Table S2.** Following the selection/isolation of these different mutations, we worked at establishing stable colonies to study each neurexin independently in a double mutant context, named here *nrnx1a1b*, *nrxn2a2b* and *nrxn3a3b*. As presented below, all these mutants and double mutants are viable. The sperm of all generated single and double mutants have been cryopreserved and are available on demand.

### Neurexins have no gross impact on the early development of zebrafish embryos and larvae

Following the successful generation of double-mutant lines, we conducted comparative phenotypic analyses. Unexpectedly, at embryonic and larval stages, none of the mutants displayed overt phenotypes in our initial comparative tests. Mutant animals hatched and developed without significant differences to their wild-type control counterparts (**Fig. 2A and 2B**). Control wild-type animals grew from 3,207 mm (± 66.9 mm) at 3 days-post-fertilization (dpf) to 3,901 mm (± 136.1 mm) at 7 dpf with no statistical differences between groups at any time point (**Fig. 2A**). All clutches tested were fully hatched by 3 dpf and grew with no observable morphological defects (**Fig. 2B**). At 2 dpf, all animals/mutants responded to touch with no difference in their swimming response/patterns compared to controls, suggesting no detectable delay or defect in brain or peripheral nervous system development at embryonic stage (**Fig. 2C**). We further analyse their behaviour up to 7 dpf using the Zebrabox Revolution platform and a battery of stimuli (**Fig. 2D and Supplemental Figure S1**). None of our experiments evidenced a significant change in their spontaneous behaviour or response to stimuli. **Figure 2D** summarises behavioural profiles of all seven double-mutant lines exposed to a 24-minute protocol combining light–dark transitions, vibration/sound, and illumination stressors. All mutants exhibited swimming patterns and response intensities comparable to controls, indicating no early obvious changes in behaviour or neural function. Finally, we monitored brain development from 2 to 7 dpf and no differences in overall structure, size, or developmental timing were observed (**Fig. 2E and Fig. 3**), supporting the conclusion that embryonic to larval nervous system development proceeds essentially normally in the absence of neurexins.

**Figure 2.**
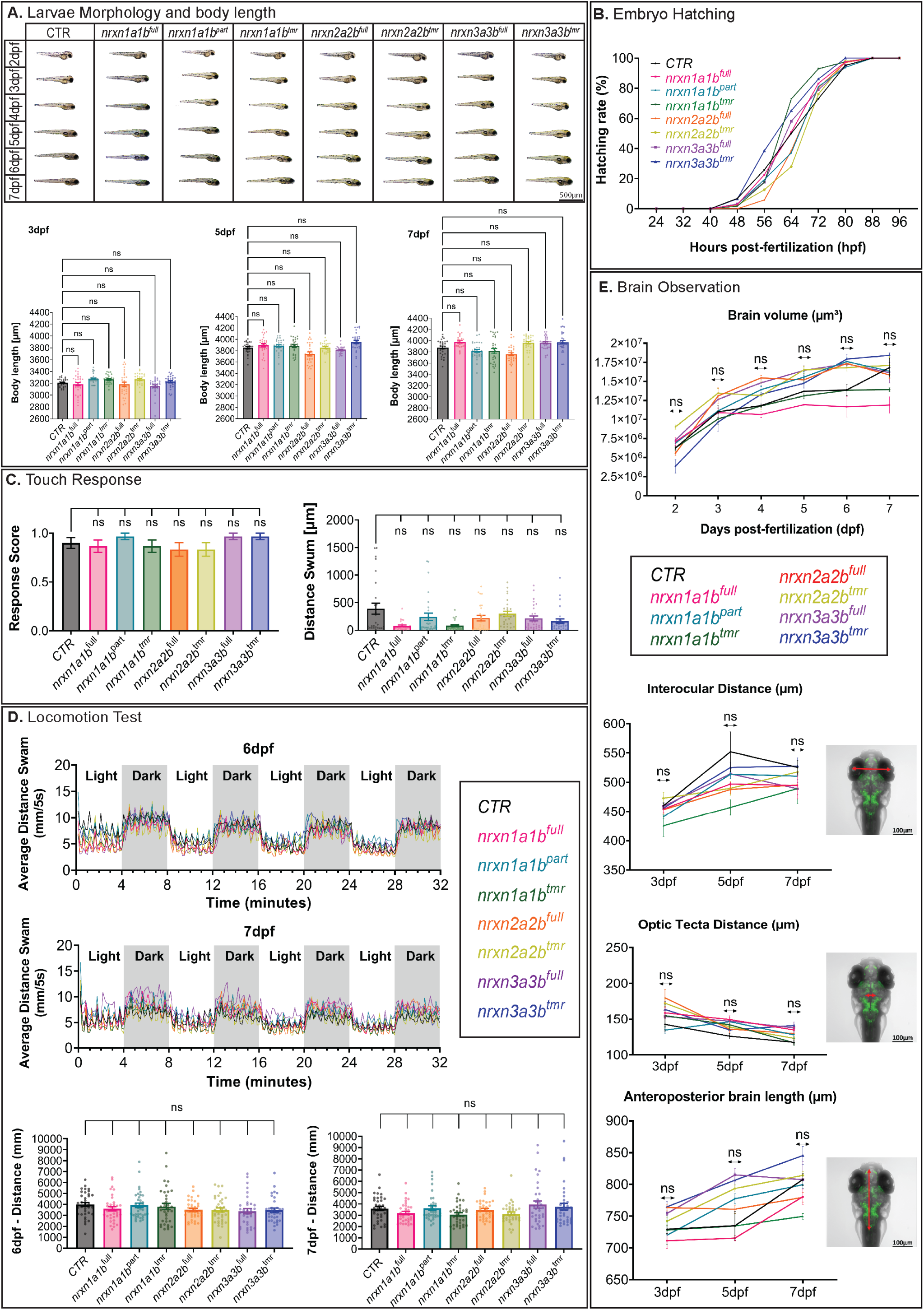
*nrxn1*, *nrxn2* and *nrxn3* are not essential to zebrafish gross development and early swimming behaviour. **A**, Upper panel, representative lateral view of the different zebrafish *neurexin* mutants versus wild-type control from 2 dpf to 7 dpf. Lower panel, body length measurement at 3, 5 and 7 dpf demonstrated no difference to wild-type control (n=30 per group; Scale bar = 500 µm. **B**, Hatching rates (%) of the *neurexin* mutants versus wild-type from 24hpf to 96hpf. **C**, Touch-evoked response scores (left panel) and swimming response (right panel) of the different neurexin mutants vs wild-type control (n=30 per group). **D**, Swimming behaviour analysis and average distance travelled during a 32-minute recording, including alternating light/dark stimulation and sound/vibration stimuli; test conducted at both 6 and 7dpf. Individual tracks are shown in **Supplemental Figure S1**. The total distance travelled across the experiment demonstrated no statistical difference across all lines tested (n=36 per group). **E**, Real-time brain development and volume analysis demonstrated no statistical difference between groups. Brain volume (upper panel) was monitored from 2 to 7 dpf, and the distance between brain landmarks (interocular/optic tectum/brainstem) was quantified at 3, 5, and 7 dpf (lower panels); Scale bar = 100 µm. Comparisons among all mutant lines and the control were performed using the Kruskal-Wallis test, followed by Dunn’s multiple comparison test. (*ns*, no significance).

**Figure 3.**
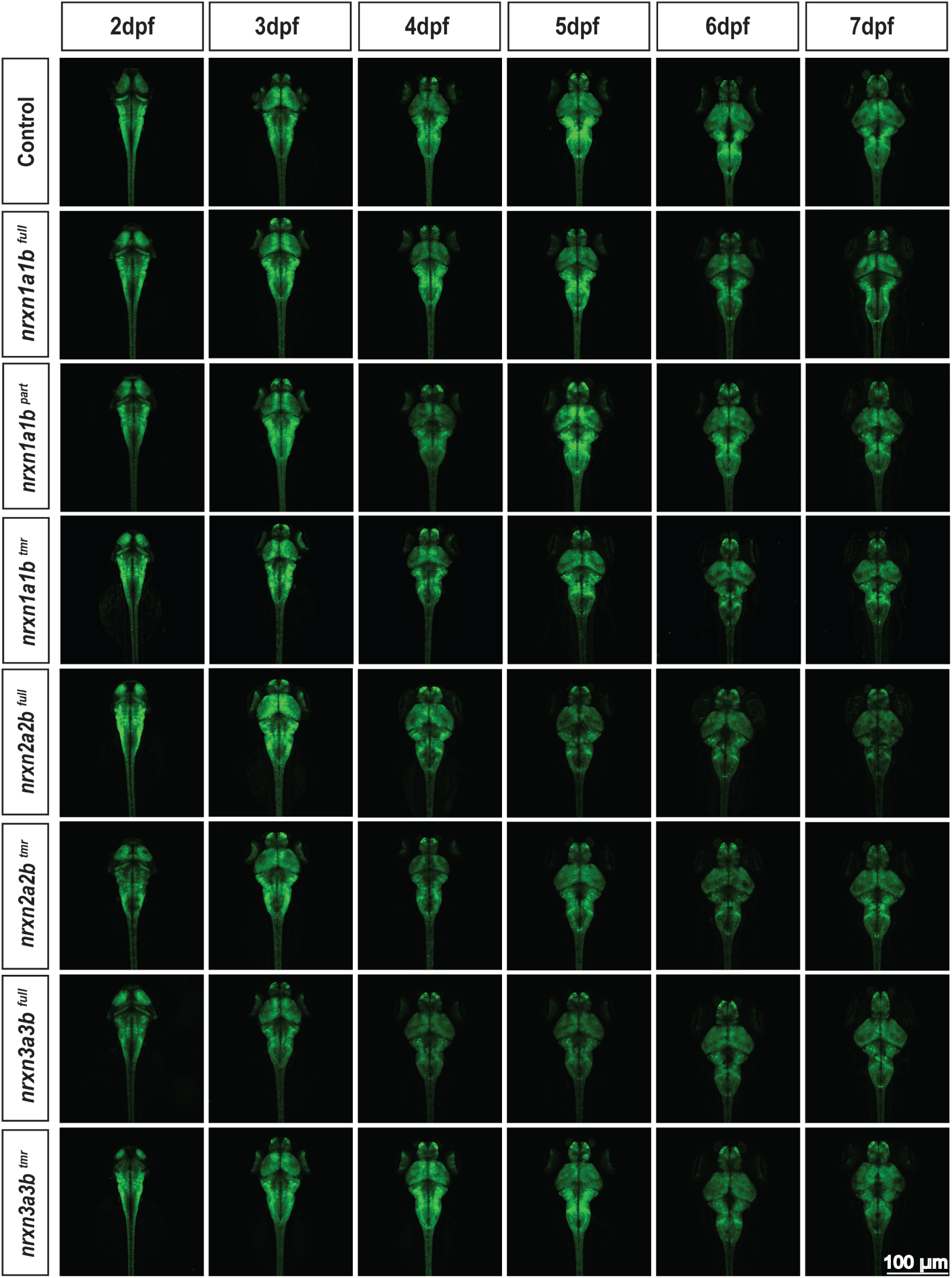
*nrxn1*, *nrxn2* and *nrxn3* are not essential for gross brain and nervous system development in zebrafish. Confocal brain scans of transgenic control (CTR; *tg(huc:Kaede)*) versus the different *neurexin* mutants in the transgenic *tg(huc:Kaede)* background. The transgene *tg(huc:Kaede)* drives expression of the fluorescent protein KAEDE in the whole nervous system. Confocal micrographs (max intensity z-stack projections) show dorsal views of whole brains acquired daily from 2 to 7 dpf, illustrating progressive brain development and the absence of overt structural abnormalities across genotypes. Scale bar = 100 µm.

### Absence of obvious early defects translated into profound adult behavioural alterations

While no obvious behavioural or developmental defects were detected at early stages, pronounced behavioural alterations emerged in juveniles to adulthood. While all mutations affected mating behaviour, rendering the maintenance and generation of embryos challenging, we observed clear paralog-specific phenotypes. This was first evident in the facility *per se.*, where *nrxn2a2b^full^* and *nrxn2a2b^tmr^* mutants displayed a nearly permanent bottom-dwelling behaviour, spending all day at the bottom of the tanks, even during feeding routines (**Supplemental Video S1**). These striking differences prompted a systematic behavioural assessment using a battery of tests, including novel tank diving, swimming endurance, social preference, mirror-biting, and shoaling assays (open-versus closed-field), presented below and in **Figures 4 to 6**.

**Figure 4.**
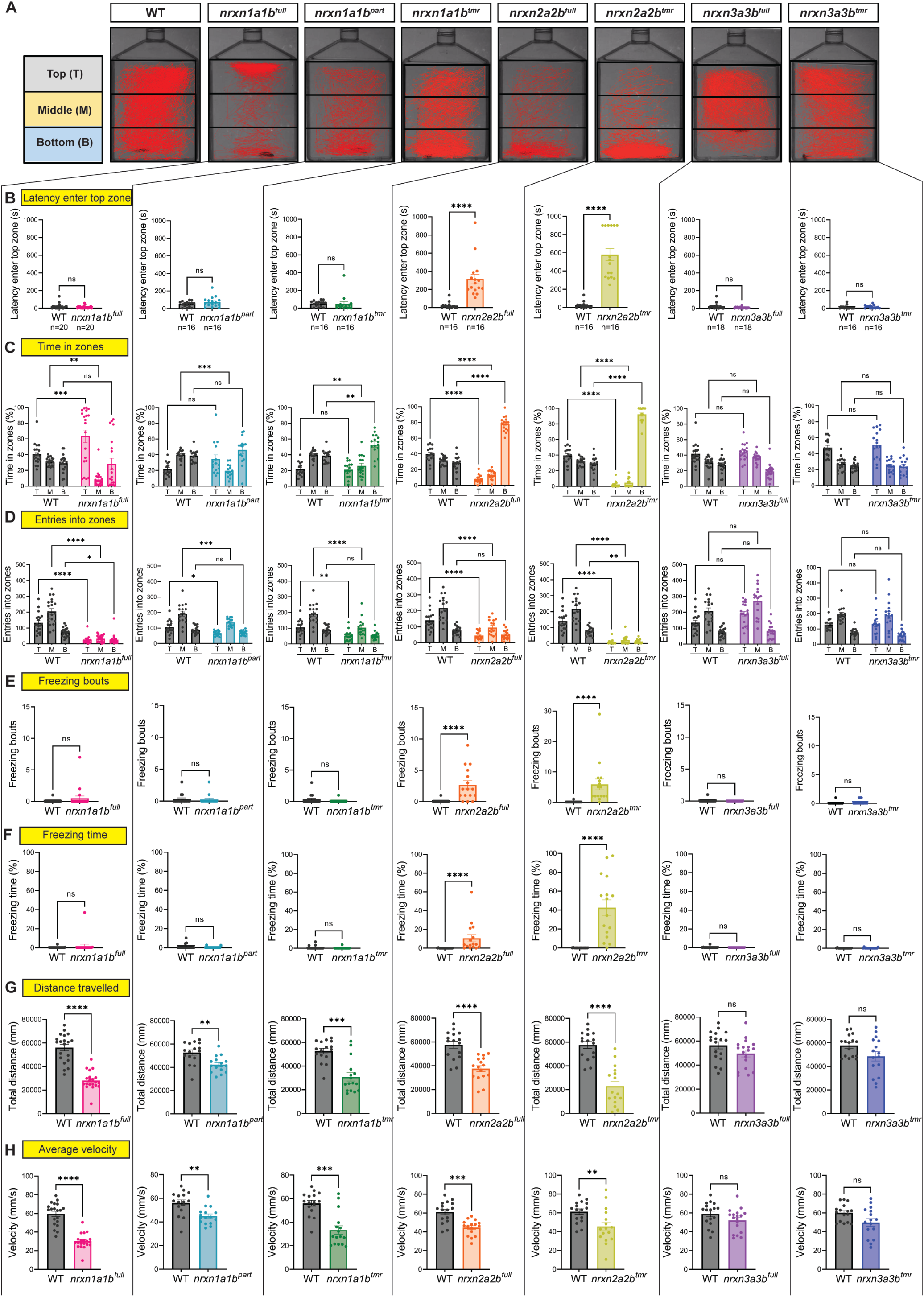
Neurexins dramatically affect adult zebrafish behaviour, with paralog-specific alterations. Novel Tank Diving Tests demonstrated that *nrxn2* deletion triggers robust anxiety-like phenotype characterised by persistent bottom-dwelling swimming coupled with repetitive freezing, whereas *nrxn1* full locus removal promotes atypical surface-swimming behaviour. A, Schematic diagram of the novel tank diving test and representative swimming tracks of wild-type versus the different *neurexin* mutant lines (15-minutes recordings). **B,** Latency to first entry into the top zone. **C,** Total time spent in top (T), middle (M), and bottom (B) zones of the tank during the 15-minute recordings. **D,** Cumulative number of entries into each zone. **E,** Cumulative number of freezing bouts. **F,** Percentage of time spent frozen during the 15-minute recording. **G,** Total distance travelled during the test. **H,** Average velocity during mobile periods (freezing periods/time excluded). Because of the large number of experimental conditions, recordings were performed across several days. Each mutant line was always tested alongside a wild-type control group recorded on the same day, ensuring that comparisons were not affected by day-to-day variability or other external influences. Data are presented as mean ± SEM. For comparisons between two groups, two-tailed Mann-Whitney tests were used. Comparisons between two groups across three zones were performed using two-way ANOVA, followed by Tukey’s multiple comparisons test. (* indicates p < 0.05, ** indicates p < 0.01, *** indicates p < 0.001, **** indicates p < 0.0001).

### *nrxn2* triggers profound anxiety-like behaviour

Following the observed consistent bottom-dwelling behaviour of *nrxn2a2b^full^*and *nrxn2a2b^tmr^* mutants within the facility, we subjected them to novel tank diving tests, a well-established assay for assessing anxiety-like behaviour in zebrafish^22^. In this paradigm, fish are introduced into a novel environment, and the latency to reach the upper section of the tank is considered inversely correlated with anxiety - *e.g.* the longer the animal stays at the bottom, the higher the stress level-. Wild-type controls reached the top section within an average of 23.16 seconds (± 8.8 seconds), whereas *nrxn2a2b^full^* and *nrxn2a2b^tmr^*mutants exhibited significantly prolonged latencies, with an average time to reach the top section of 314.4 seconds (± 8.8 seconds) (p < 0.0001) and 580.1 seconds (± 66.8 seconds) (p < 0.0001), respectively (**Fig. 4A and 4B**). Strikingly, 37.5% of *nrxn2a2b^tmr^* mutants never explored up to the upper section, remaining exclusively at the bottom throughout the assay, indicating a severe anxiety-like state in zebrafish.

Consistent with these findings, analysis of vertical distribution and number of transitions between the three zones revealed a marked reduction of their exploratory behaviour when compared to controls (**Fig. 4C** and 4**D**). While wt fish spent a cumulative 28.6% (± 2.3%) of the assay in the lower section, *nrxn2a2b^full^* and *nrxn2a2b^tmr^* mutants remained almost exclusively at the bottom, spending 78% (± 2.4%) (p < 0.0001) and 92% (± 2.4%) (p < 0.0001) of the total experimental time in the lower section, respectively (**Fig. 4C**). Demonstrating not only a bottom-dwelling but also a clear reduction of their explorative behaviours, while wild-type fish performed 421 (± 45) cumulative transitions between sections during the assay, *nrxn2a2b^full^* mutants made only 192 (± 25) (p = 0.0003) transitions, and *nrxn2a2b^tmr^* mutants rarely exited the bottom zone, with only 43 (± 15) (p < 0.0001) total transitions across the trial (**Fig. 4D**).

### *nrxn2* triggers robust and repetitive freezing/seizures, but does not affect swimming performance

In addition to a profound bottom-dwelling and reduction of exploration during our test, *nrxn2a2b^full^* and *nrxn2a2b^tmr^*also displayed repetitive and robust freezing/seizure behaviours often followed by a burst of activity, both during behavioural assays and under routine facility observation (**Fig. 4E and 4F, Supplemental Video S2**). In the novel tank diving tests, *nrxn2a2b^tmr^* displayed freezing behaviour for a mean of 43% (± 8%) of the experimental time. By contrast, wt controls showed no freezing (mean 0.03%). In comparison, *nrxn2a2b^full^*froze for 11% (± 4%) of the experimental time, appearing less affected than *nrxn2a2b^tmr^*. In addition, *nrxn2a2b* mutants, along with all *nrxn1a1b* but not *nrxn3a3b* mutants, showed a marked reduction in total distance travelled and average swimming velocity during these novel tank diving experiments (**Fig. 4G and 4H)**, these values were calculated after excluding freezing periods to avoid bias). To determine whether these reductions reflected neuromuscular/fitness impairment rather than altered behaviour, the different lines were subjected to an endurance assay using a Loligo® swimming tunnel, in which the animals were challenged with a progressively increasing water flow. As presented in **Fig. 5A**, mutants and wt animals displayed comparable resistance to increasing flow, indicating preserved strength and endurance, suggesting that the observed reduced distance swam and velocity arise primarily from behavioural alterations rather than motor deficits.

**Figure 5.**
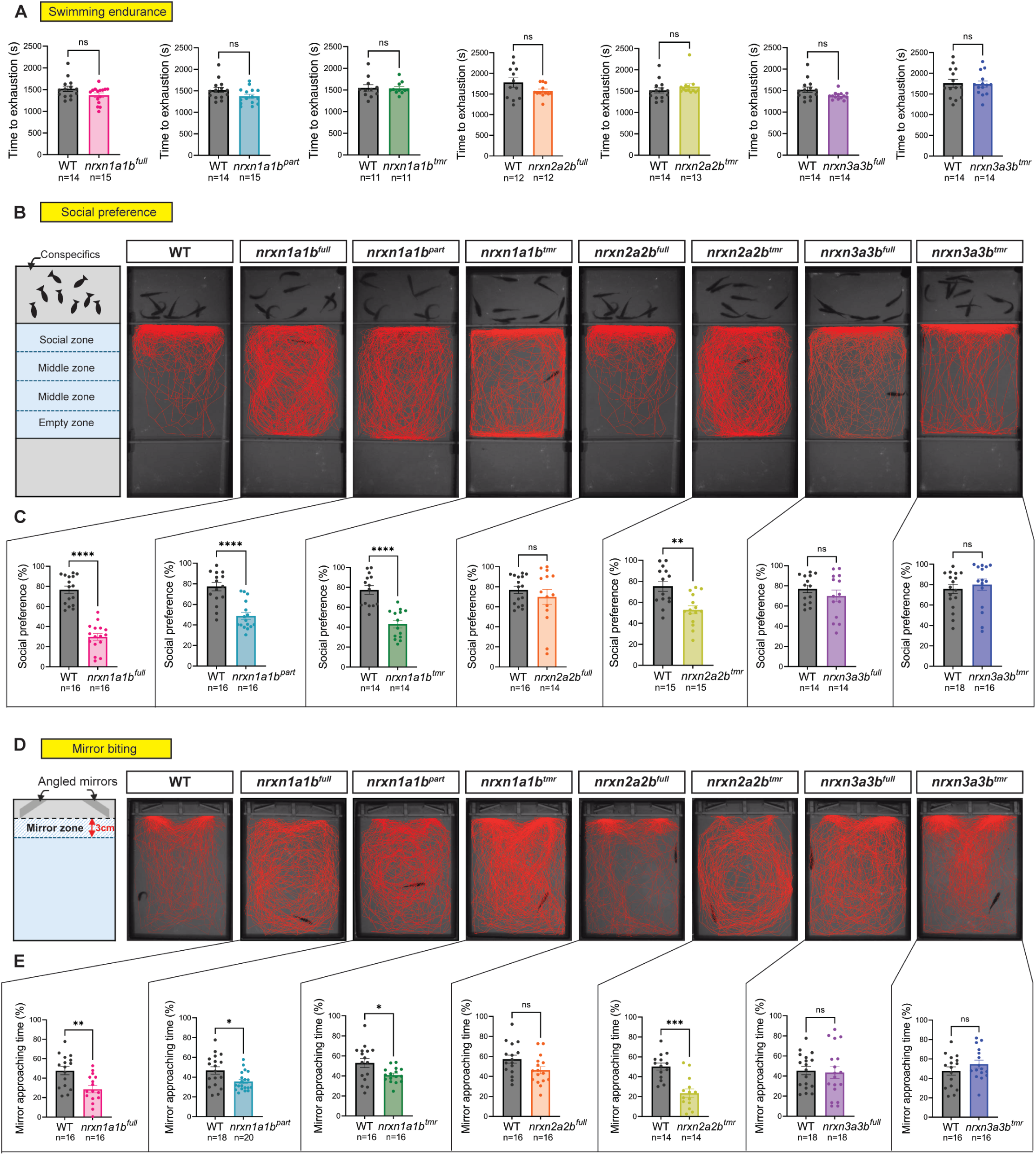
*nrxn1* modulates social behaviour and aggressivity in zebrafish. **A**, Swimming endurance assays, with performance expressed as time to exhaustion. **B-C,** Social preference assays with: **B,** Schematic diagram of the social preference test/apparatus, and representative swimming tracks of wild-type versus the seven *neurexin* mutant lines during 10-minute recordings. **C,** Quantification of the percentage of time the tested fish spent in the social zone. **D-E,** Mirror biting assays with: **D,** Schematic diagram of the mirror biting test/apparatus and representative swimming tracks of wild-type versus the different *neurexin* lines during the 10-minute test. **E,** Percentage of time the fish spent in the mirror zone. Recordings were performed across several days, with each mutant tested alongside same-day wild-type controls to avoid day-to-day variability. Data are shown as mean ± SEM. For comparisons between two groups, two-tailed Mann-Whitney tests were used. (* indicates p < 0.05, ** indicates p < 0.01, *** indicates p < 0.001, **** indicates p < 0.0001).

### *nrxn1* triggers unusual surface-dwelling behaviour coupled with reduced exploration

In contrast to *nrxn2a2b*, *nrxn1a1b* mutants did not exhibit strong anxiety phenotypes, but instead demonstrate marked defects/changes in their social behaviours and aggressivity (see below and **Fig. 5**). Before describing these new sets of assays/phenotypes, it is noteworthy to highlight that, during the novel diving tests, none of the *nrxn1a1b* mutants demonstrated significant change in their latency to enter the top zone (**Fig. 4A and 4B** – concluding on an absence of significant anxiety). However, *nrxn1a1b^full^,* specifically, demonstrated a pronouncedly significant top-section swimming behaviour/preference (**Fig. 4C**), combined with less exploration/transition to the middle and bottom zones and less overall distance travelled (**Fig. 4D, G and H, Supplemental Video S2**). To our knowledge, this combination of marked top section preference coupled with reduced exploration and reduced distance-swam/velocity has not been previously described and suggests profound and potentially complex behavioural alteration. Interestingly, while *nrxn1a1b^part^* and *nrxn1a1b^tmr^* did not share this significant top section preference in the novel tank diving test, they demonstrated a similar significant surface-dwelling behaviour in the closed field tests presented below (**Fig. 6**).

### *nrxn1* modulates social behaviour and aggressivity

In addition to their surface-dwelling behaviour, all *nrxn1a1b* mutants developed marked changes in social behaviour and aggressivity (**Fig. 5B-E, Supplemental Video S3 and Supplemental Video S4**). To assess social behaviour, we used three-chambered experimental tanks in which two small side compartments were separated from the central arena by transparent glass. One side compartment contained a group of conspecifics, while the opposite compartment was left empty. The test fish was placed in the central arena, and its spatial distribution was analysed to quantify social preference/behaviour. As shown in **Fig. 5B and 5C**, while wild-type control spent 77% (± 4%) of the experimental time adjacent to their sibling, *nrxn1a1b^full^*, *nrxn1a1b^part^* and *nrxn1a1b^tmr^* mutants exhibited a pronounced social withdrawal, spending only 30% (± 3.4%), 49% (± 3.5%) and 43% (± 3.8%) of the experimental time, respectively, near conspecifics. To assess aggressiveness, we deployed a mirror biting test (**Fig. 5D and 5E**). Wild-type control animals spent an average of 48% (± 3.7%) within the 3cm zone next to the mirror. In contrast, all *nrxn1a1b* mutants demonstrated a significant reduction of their time interacting with the mirror, with 29% (± 3.7%) for *nrxn1a1b^full^*, 36% (± 2%) for *nrxn1a1b^part^*and 40% (± 1.9%) for *nrxn1a1b^tmr^*. To further support these results, the *nrxn1a1b^full^* mutants we further subjected to a dyadic fighting test^23^. As presented in **Supplemental Fig. S3**, the mirror-biting findings were corroborated by a robust decrease in direct aggressive interactions, with mutants displaying only 2 (± 1) bites per minutes on average, compared with 16 (± 3) bites per minute in the wild-type controls (p < 0.05). It is noteworthy that the mirror biting tests also concluded on a significant alteration of social behaviour and aggressivity for *nrxn2a2b^tmr^*animals; however, the high frequency of freezing and seizure-like episodes observed during the tests may have confounded these results, and the significance of these findings should therefore be interpreted with caution.

### *nrxn1* and *nrxn2* display opposite responses to open and closed field tests

Building on the initial behavioural analyses conducted with single animals, we next assessed group behaviour by analysing shoal distribution in an open versus closed field apparatus. The main goal was to confirm an observation made in the facility during daily maintenance; *nrxn2a2b* mutants, typically exhibiting persistent bottom-dwelling, began to explore their tanks when placed in a protected or covered area. This contrasted sharply with their constant bottom-dwelling behaviour when tanks were positioned on the open workbench or openly exposed in the facility racks. For the experiment, the same test tanks were used in both open- and closed-field conditions; the key distinction was that in the closed-field setup, the tank was fully covered, providing shelter and shielding the fish from external cues (**Fig. 6B**). In contrast, in the open-field setup, the tank remained uncovered, leaving animals fully exposed to their surroundings. Groups of ten fish were introduced in the testing tanks, and fish distribution was analysed over time and plotted in **Figure 6**. Replicating the observations made in the facility and during the novel tank diving test, all *nrxn2a2b* mutants in the open field apparatus displayed a robust and persistent bottom-dwelling behaviour, with an average of 80% (± 4%) and 99% (± 0.3%) of the shoal of *nrxn2a2b^full^*and *nrxn2a2b^tmr^* distributed at the bottom of the tanks across the entire duration of the experiment; defects highly significant compared to wild-type controls, with only 17% (± 6%) of their shoal recorded at the bottom compartment across the experiment (p < 0.0001) (**Fig. 6I and 6K**). Confirming our observation, when transferred to closed-field conditions, *nrxn2a2b^full^*did not behave differently from wild-type, with the shoal distribution in each zone being the same as in wt controls (**Fig. 6J**). *nrxn2a2b^tmr^* mutants still differed from wt controls but showed clear improvement, with bottom-dwelling reduced from 99% (± 0.3%) in open field to 68% (± 5.5%) when transferred to closed field.

**Figure 6.**
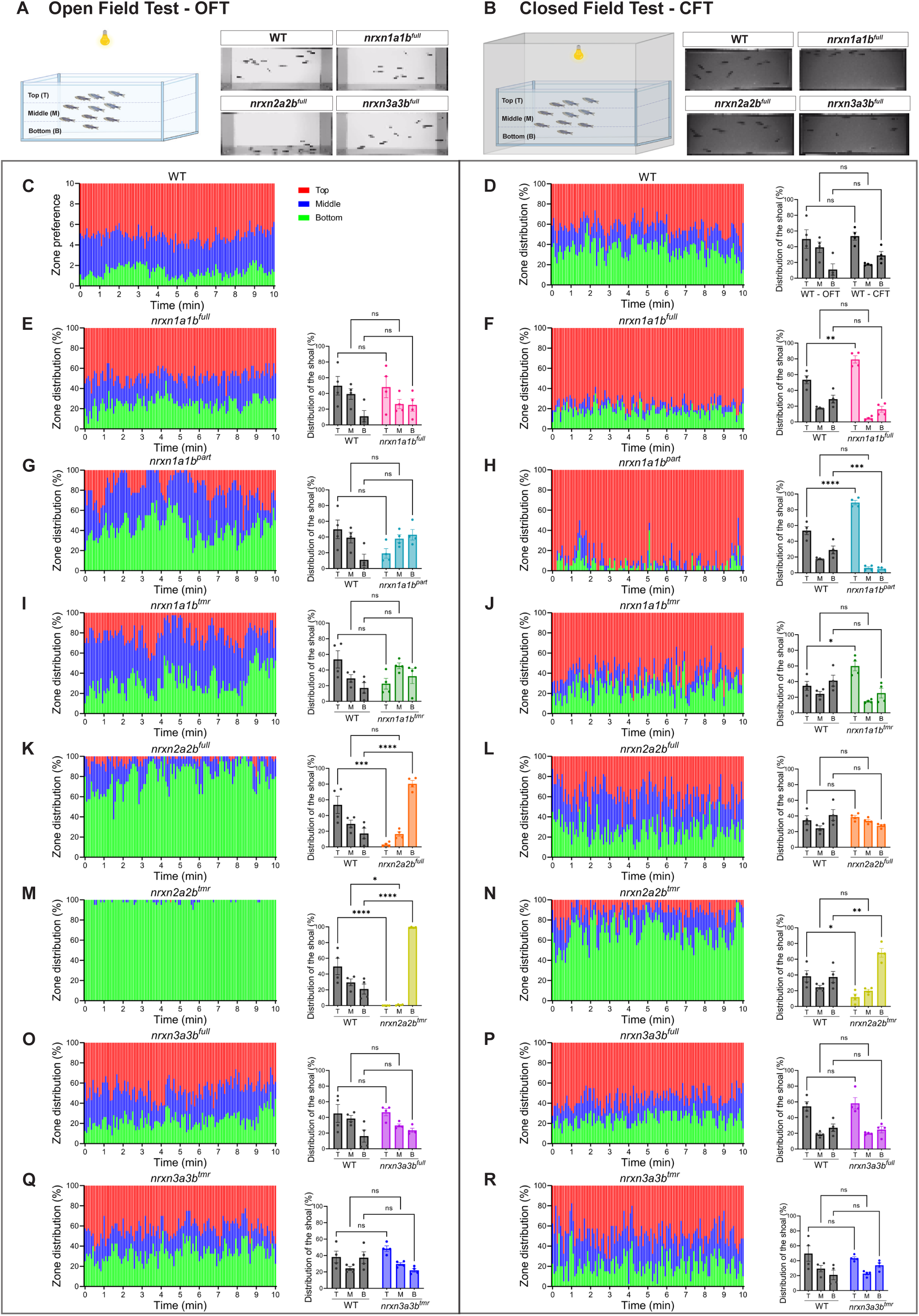
*nrxn1* and *nrxn2* mutant lines display a divergent response to open and closed field environments. **A**, Schematic representation of the Open Field Test apparatus and video-snapshots examples. **B,** Schematic representation of the Closed Field Test apparatus and video-snapshots examples. **C-R,** Zone preference over time of wild-type and the seven studied neurexin mutants. **Left panels** represent the average number of fish in each zone every 5s through the 10 minutes recordings (Percentage of 10 fish per 5s, with 4 independent groups; n=4 with 40 fish total). **Right Panels** average the overall spatial distribution of the 10 recorded fish over the total experimental recording of 10 minutes (Percentage of 10 fish over 10 minutes recordings, with 4 independent groups; n=4 with 40 fish total). Experiments started after 5 minutes of habituation. Recordings were performed across several days, with each mutant tested alongside same-day wild-type controls to avoid day-to-day variability. Data are shown as mean ± SEM. For comparisons between two groups across three zones, two-way ANOVA was applied, followed by Tukey’s multiple comparisons test (* indicates p < 0.05, ** indicates p < 0.01, *** indicates p < 0.001, **** indicates p < 0.0001).

In contrast, while transition to a closed/covered environment did not impact wt or *nrxn3a3b* mutants’ behaviour (**Fig 6C-D, 6M-P**) and rescued/reduced *nrxn2a2b* defects, it did trigger significant alteration of all *nrxn1a1b* mutants’ behaviour (**Fig. 6C-H**). While all *nrxn1a1b* mutants behave within wild-type control ranges in the open field set-up, transfer to closed-field conditions triggered a significant surface-dwelling behaviour and reduction of exploration, with *nrxn1a1b^full^*, *nrxn1a1b^part^* and *nrxn1a1b^tmr^* mutants spending 79% (± 4.6%, p < 0.01), 88% (± 3%, p < 0.0001) and 60% (± 6%, p < 0.05) of the experimental time swimming at the surface. To our knowledge, this constitutes an undescribed behavioural response in such a paradigm, suggestive of profound and complex circuitry alterations, and may be particularly relevant given the strong association of human *NRXN1* mutations with schizophrenia, a condition characterised by altered sensorimotor gating and abnormal processing of environmental context.

In short, open to closed field transition did not affect wild-type or *nrxn3a3b* mutants’ behaviour but significantly rescued or reduced *nrxn2a2b*-induced anxiety-like behaviour, while, oppositely, significantly aggravated or triggered *nrxn1a1b* altered surface-dwelling behaviour.

### *nrxn3* does not trigger obvious abnormal change from early to adult stage

By comparison, *nrxn3a3b* double mutants developed normally and did not display overt alterations in exploratory, anxiety-like, or social behaviour (**Fig. 4, 5 and 6**). Across novel tank diving, mirror-bitting, social interaction and open/closed shoal field tests. all *nrxn3a3b* mutants consistently performed within control ranges. These results indicate that *NRXN3* is either dispensable for the measured adult behaviours or functionally compensated in the studied mutants.

## DISCUSSION

In this study, we generated the first complete collection of zebrafish neurexin knockout lines and the first comparative phenotypic analysis, which underscores the distinct and non-redundant roles of neurexin paralogs in shaping juvenile/adult behaviours. Interestingly, the absence of early obvious developmental defects, combined with the emergence of pronounced adult behavioural alterations, resembles the clinical trajectory of many neurodevelopmental and psychiatric disorders such as schizophrenia, autism, and bipolar disorder, which often lack overt neuroanatomical hallmarks yet manifest with strong and rapidly progressing behavioural dysfunctions^1,24^. These findings highlight both the conserved and complex role of neurexins in brain function as well as the utility of zebrafish as a complementary model system to dissect the genetic and circuit-level mechanisms underlying psychiatric conditions. The extreme and robust phenotypes observed with *nrxn1* and *nrxn2* LOF lines clearly exemplify the potential of the zebrafish for studying the role of genetics in mental health and behavioural disorders.

Our data show that *nrxn1* and *nrxn2*, but not *nrxn3*, are critical for the regulation of social behaviour, anxiety, and aggression in zebrafish. Loss of *nrxn2* produced a striking and penetrant anxiety-like phenotype, characterised by bottom-dwelling, reduced exploration, and frequent freezing/seizure-like episodes. These findings resonate with rodent studies implicating *Nrxn2* in anxiety-related behaviours and with human genetic associations linking *NRXN2* variants to autism and intellectual disability^4,25^. Similarly, the robust and repetitive freezing observed in *nrxn2* mutants, particularly the spontaneous mid-water freezing noted in the facility outside of induced stress paradigms, warrants further investigation. These episodes strongly resemble seizure/epileptic-like activity rather than the fear-related freezing typically described in zebrafish novel tank diving tests. Notably, during such events, animals did not respond to manipulation or attempt to evade nets or handling tools, which deviates from classic defensive freezing responses. Clarifying whether these episodes represent epileptiform activity or an extreme anxiety response will require *in vivo* imaging or electrophysiological validation. This observation is especially relevant given the growing links between *NRXN2*, epilepsy, and neurodevelopmental disorders in patients^25,26^. Conversely, *nrxn1* deletion did not elicit strong anxiety-like behaviour in zebrafish but instead induced profound alterations in social behaviour, aggression, and an unusual surface-dwelling phenotype that was exacerbated under closed-field conditions. These results resonate with the fact that *NRXN1* has been strongly associated with schizophrenia, a disorder characterised by profound social dysfunction and, in many patients, increased aggression^1,4,27^.

It is noteworthy that these phenotype divergences between *nrxn1* and *nrxn2* mutants are of particular interest. Confinement alleviated anxiety-like behaviours in *nrxn2* mutants but paradoxically triggered aberrant surface-dwelling in *nrxn1* mutants. This suggests that different neurexin paralogs gate how environmental cues are integrated into behavioural output. As briefly stated above, in humans, *NRXN1* deletions have been strongly linked with schizophrenia and abnormal sensory gating, while *NRXN2* has been linked with heightened anxiety and ASD-like features. Our zebrafish data therefore also support the notion that *nrxn1* and *nrxn2* act through distinct neural pathways that differentially process environmental and contextual information, which could explain why different paralogs are associated with different psychiatric conditions.

Another striking observation is that none of the zebrafish *nrxn* mutants displayed gross developmental or morphological defects during embryonic or larval stages. Brain volume, gross anatomy, and basic locomotor responses were all indistinguishable from wild-type controls. Yet, profound phenotypes emerged later in juvenile to adult stages. This mirrors the clinical paradox of psychiatric disorders, which are widely recognised to have early neurodevelopmental origins yet lack obvious developmental abnormalities. This “silent developmental trajectory” supports the hypothesis that neurexin dysfunction perturbs synaptic wiring or plasticity in ways that remain latent until circuits mature or environmental and social demands increase. Zebrafish, with their optical accessibility and powerful *in vivo* imaging tools early in life, are ideally suited to investigate the precise timing and mechanisms of such latent/subtle circuit disruptions.

By contrast, *nrxn3* mutants did not display obvious abnormalities in the behavioural assays performed, but this does not exclude a possible effect on behavioural domains not captured by our paradigms.

## CONCLUSION

In summary, our study introduces the first complete set of zebrafish mutants spanning the entire *neurexin* gene family. Despite showing no overt neurodevelopmental or motor defects during embryonic and larval stages, these lines reveal pronounced and paralog-specific behavioural disturbances that emerge only during juvenile-to-adult transitions. *nrxn1* loss in zebrafish primarily alters social behaviour and aggressivity, while *nrxn2* triggers severe anxiety-like behaviours and freezing/seizure-like events. *nrxn3* loss, however, appears dispensable in all domains tested, which is consistent with a less robust association between *NRXN3* and mental health disorders in humans.

These phenotypes trajectories in zebrafish mirrors a central challenge in human mental health research: most psychiatric and neurodevelopmental disorders are widely accepted to originate early in life, yet often present without clear, detectable abnormalities during early development. By uncovering dramatic adult phenotypes in the absence of obvious early defects, these mutants create a unique opportunity to explore the subtle, progressive, and circuit-level mechanisms that unfold between early development and behavioural manifestation. The optical accessibility, genetic tractability, and whole-brain imaging capacity of the zebrafish make it an exceptionally powerful model to dissect how early molecular perturbations in synaptic adhesion genes translate into late-onset behavioural pathology. As such, these *nrxn* mutants open a new avenue to investigate the developmental timing, network rewiring, and mechanistic origins of mental disorders, with direct relevance for understanding disease onset and for identifying novel therapeutic windows.

As a whole, this study also demonstrates the tremendous value of the zebrafish as a model to investigate the underlying role of genetics in mental illnesses.

## Supporting information

Supplemental File S1 (Sequences)

## ACKNOWLEDGMENT

We would like to thank the Queensland Centre for Mental Health Research and Queensland Health for their long-term support in the generation of the different neurexin lines. The generation of these mutants is very time-consuming and would not have been possible without their support.

## FUNDING

This work was supported by grants from the Australian National Health and Medical Research Council (NHMRC) Project Grant No 1165850 to B.M. and J.G., the Rebecca Cooper Medical Research Project Grant No PG2019405 to J.G. and the NHMRC Emerging Leader Fellowship No 1174145 to JG.

## CONFLICT OF INTEREST

None to declare

## Notes

### Competing Interest Statement

The authors have declared no competing interest.

