## Supplemental Material for "Modelling mental disorders in zebrafish. Neurexins severely modulate anxiety, social behaviours and aggression"

**Table of contents:**

**Supplemental Table S1.**	Page 2

**Supplemental Table S2.**	Page 3

**Supplemental Figure S1.**	Page 4

**Supplemental Figure S2.**	Page 5

**Supplemental figure S3.**	Page 6

**Supplemental Video S1.**	Page 7

**Supplemental Video S2.**	Page 7

**Supplemental Video S3.**	Page 7

**Supplemental Video S4.**	Page 7

**Supplemental File S1.**	Page 7

**Supplemental Table S1.** **CRISPR/Cas9 guide RNAs and oligonucleotides used in this study.**

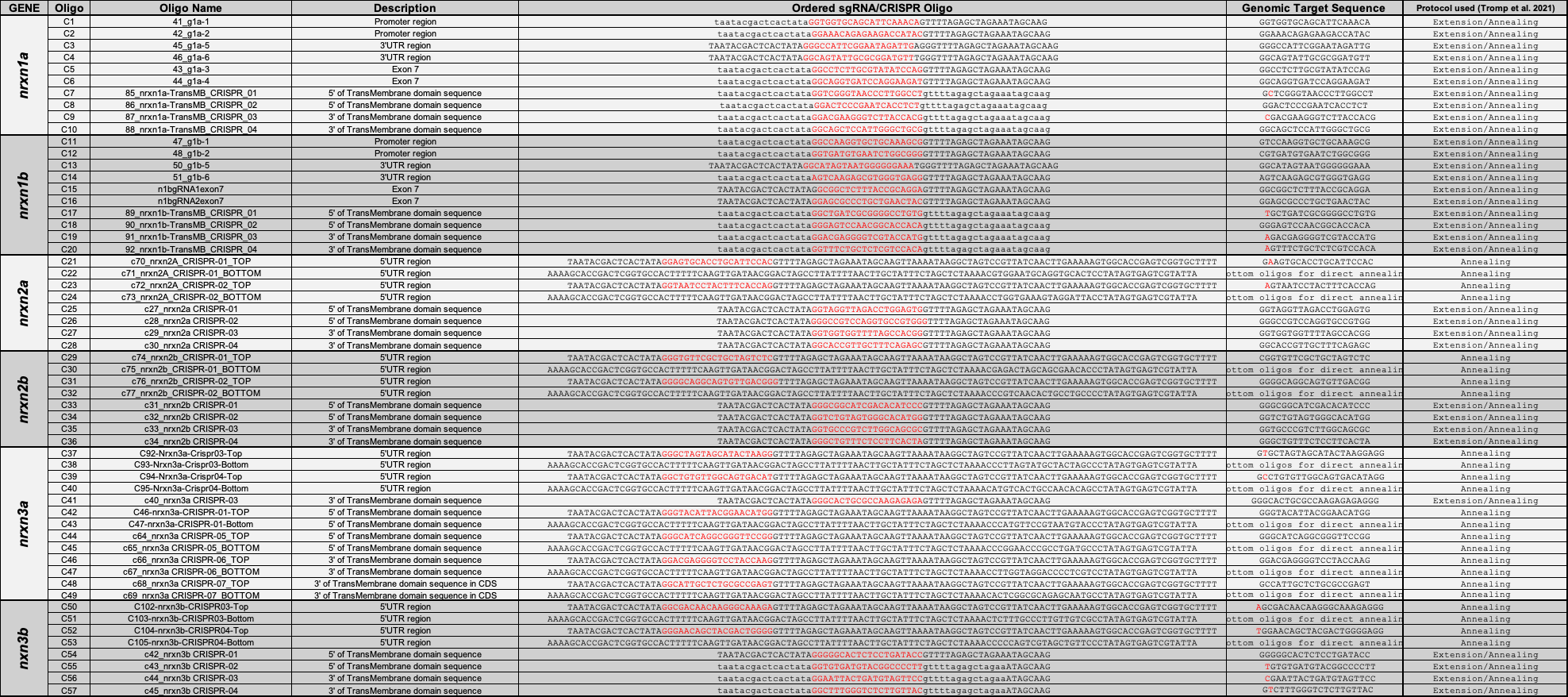

**Supplemental Table S2. NRXN genotyping primers used in this study.**

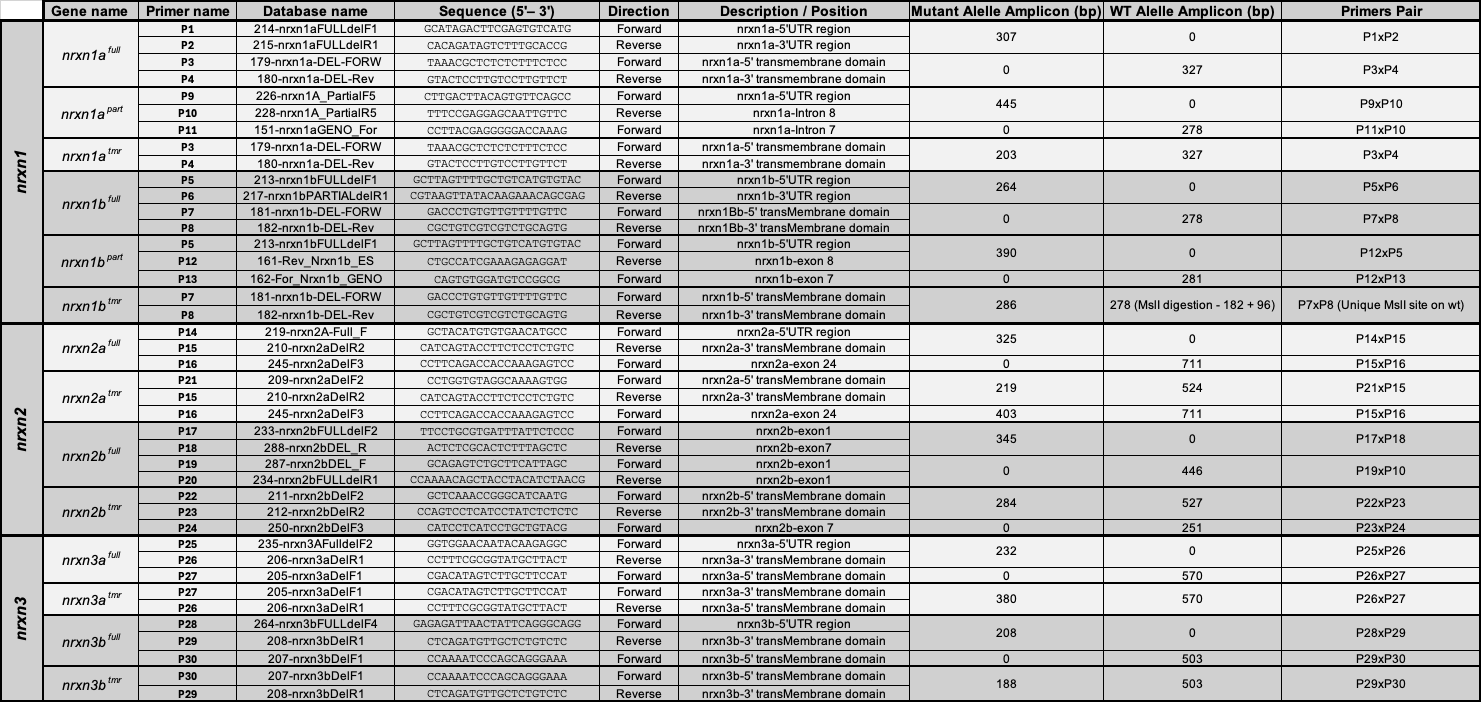

**Supplemental Figure S1. Larval swimming patterns and locomotor activity of wild-type controls versus the different neurexin mutant lines presented in this study.** Total distance moved (mm) of 6 dpf larvae (plotted per 5-second bin) over 32-minute recordings. No significant differences were observed between mutants or controls.

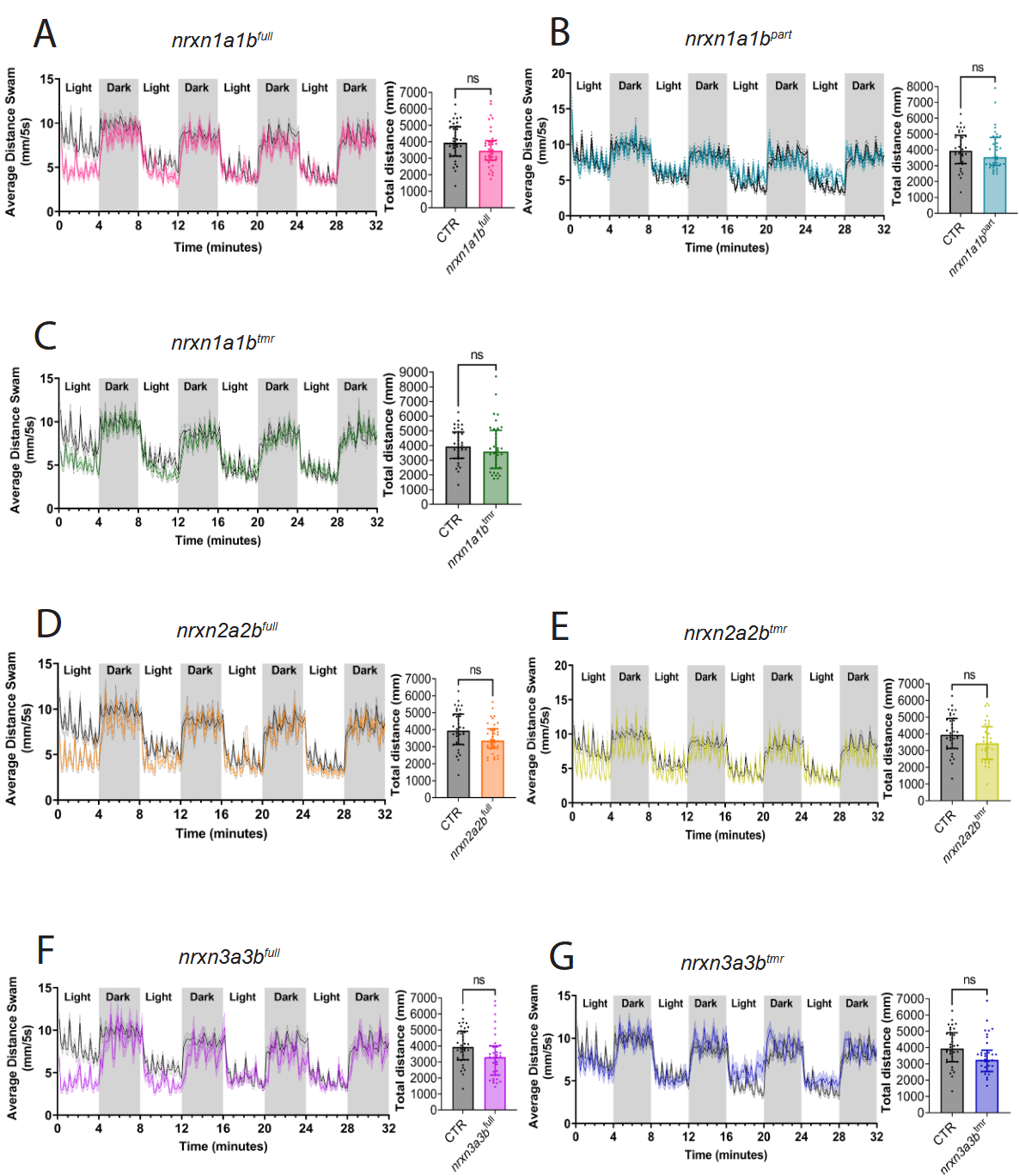

**Supplemental Figure S2. Larval swimming patterns and locomotor activity of wild-type controls versus the different neurexin mutant lines presented in this study.** Total distance moved (mm) of 7 dpf larvae (plotted per 5-second bin) over 32-minute recordings. No significant differences were observed between mutants or controls.

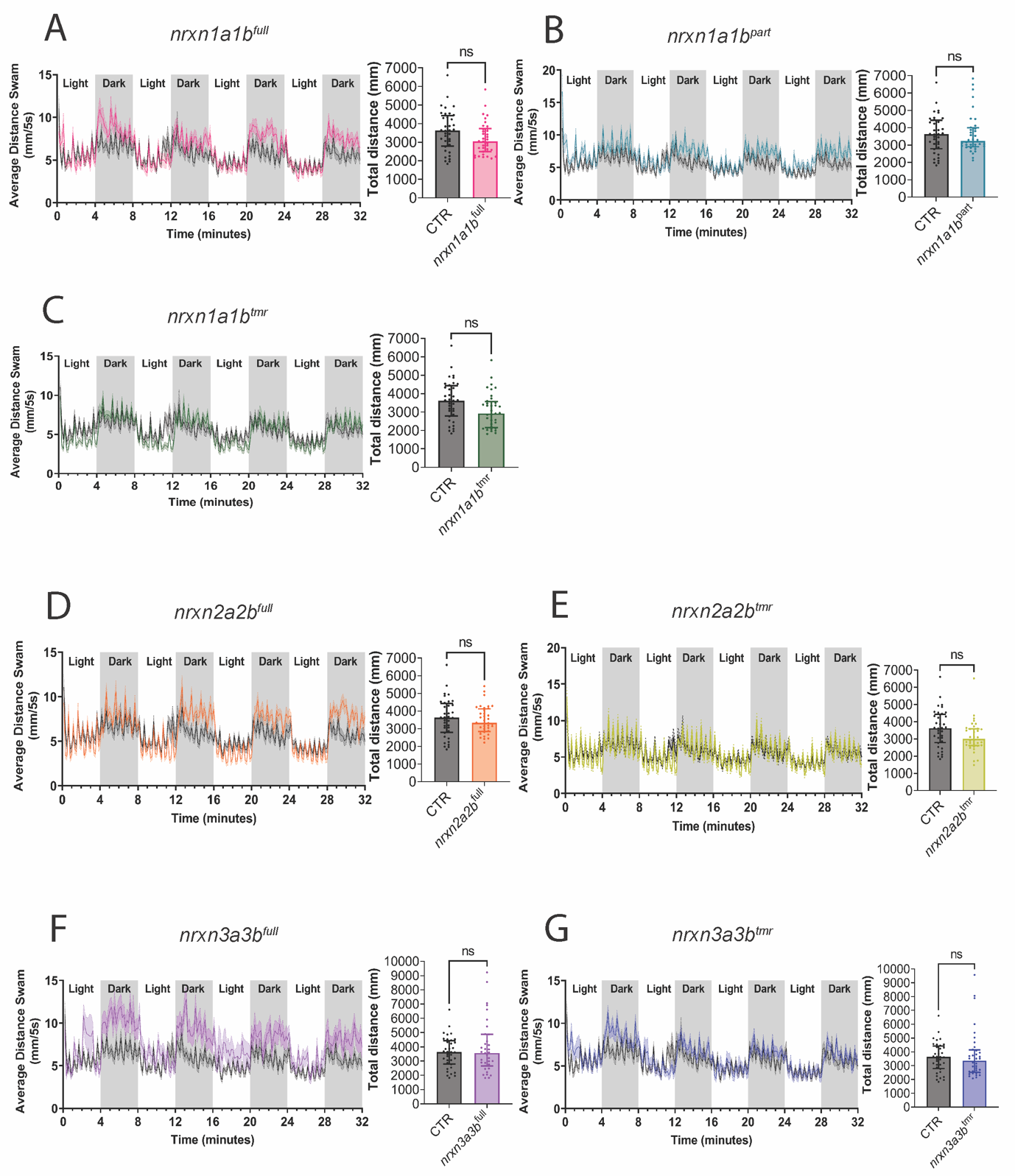

**Supplemental figure S3. *nrxn1* locus deletion results in reduced aggressiveness (dyadic fighting test).** Two size-match male fish of each genotype group were placed in a behavioral tank separated by an opaque divider and acclimated for 24h before the test to induce aggressiveness. After the 24-h acclimation, the divider was removed, and the fish were allowed to interact freely for 30 min. The number of bites was quantified using the manual scoring function of EthoVision XT software (Ver. 18.0.1803, Noldus) and plotted using GraphPad Prism (version 9.0.0). **A**, Number of bites recorded per minute during the 30-minute interaction. **B**, Average number of bites over the entire 30-minute recording period (n=4; 2 fish per point). Data are shown as mean ± SEM. Comparisons between two groups were performed using a two-tailed Mann-Whitney test (p < 0.05).

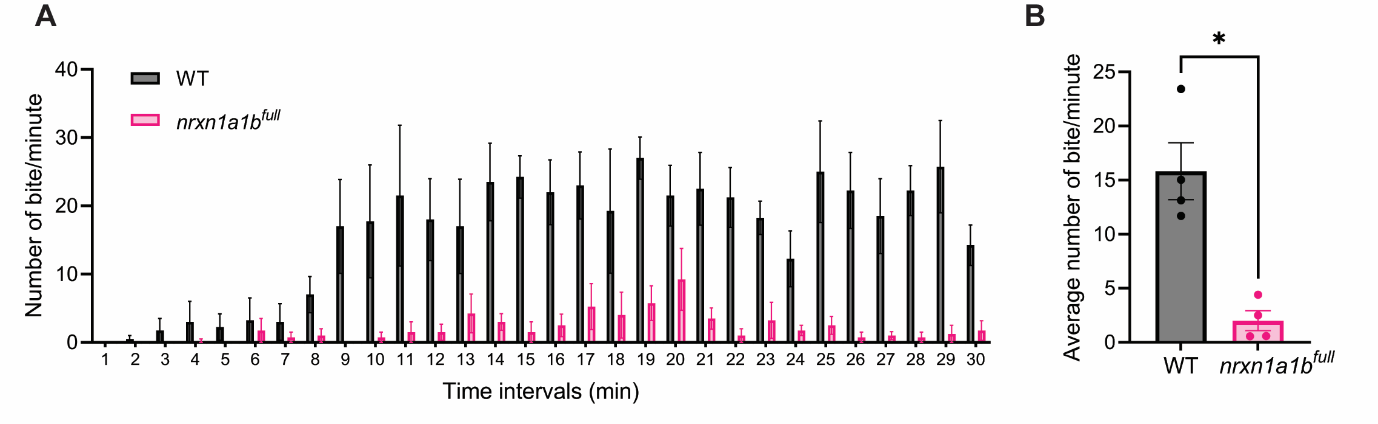

**Supplemental Video S1. rxn2 mutants exhibit a near-constant bottom-dwelling behaviour under standard facility conditions.** This 4-minute recording shows the different neurexin lines within their respective home tanks, filmed immediately after removal from the facility rack.

*Available online to download*

**Supplemental Video S2. Representative (4× speed) video recordings of the Novel Tank Diving Tests performed in this study.**

*Available online to download*

**Supplemental Video S3. Representative video recordings of the Social Behavior assays conducted in this study.**

*Available online to download*

**Supplemental Video S4. Representative video recordings of the Mirror Biting assays conducted in this study.**

*Available online to download*

**Supplemental Material S1. Zip file containing snapgene files of the different neurexin zebrafish genes with guides, primers and deletions generated in this study.**

*Available online to download*
